# Improving interpretability of transcription factor binding models with DNA shape features

**DOI:** 10.1101/2025.04.01.646034

**Authors:** Ryan L. Keivanfar, Forest Yang, Katherine S. Pollard, Nilah M. Ioannidis

## Abstract

Deep learning models in genomics that predict molecular phenotypes from DNA sequence traditionally focus on one-hot encoded nucleotide representations. Here, we develop a novel model that extends this approach by incorporating DNA structural attributes indicative of local DNA shape alongside canonical sequence inputs. This augmentation provides an additional axis for model interpretability and aids in identifying regulatory patterns not apparent from sequence alone. Applying this approach to prediction of transcription factor binding (ChIP-seq) demonstrates that combining sequence and structural DNA information can improve the identification of regulatory elements to provide a more nuanced understanding of genomic function and regulation.

**Graphical abstract:** 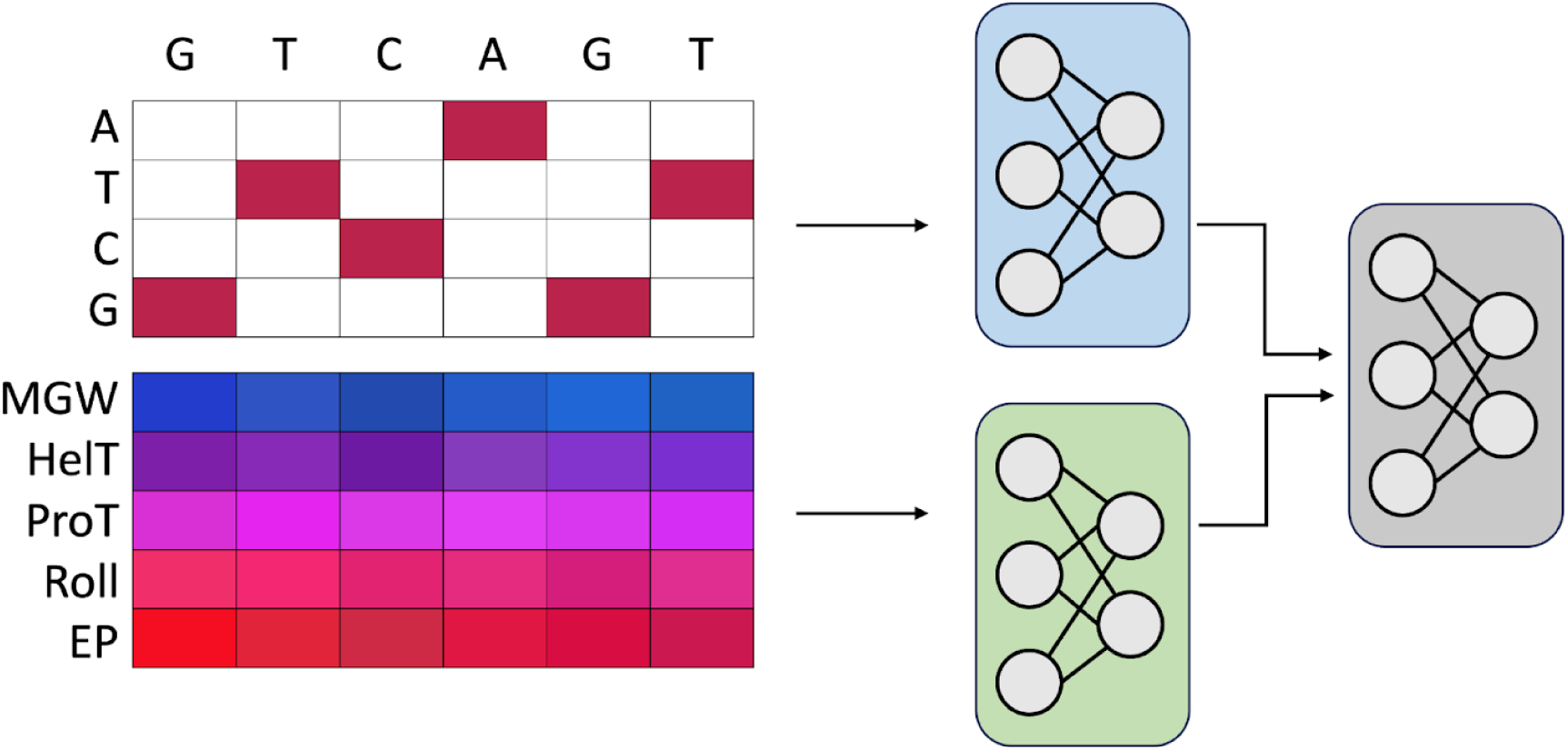

Schematic overview of the DeepShape model. One-hot encoded sequence, and five DNA shape attributes—minor groove width (MGW), helical twist (HelT), propeller twist (ProT), roll, and electrostatic potential (EP)—are input separately to sequence and shape branches of the model, each processed through two convolutional layers and subsequently concatenated for further processing through additional convolutional layers.

## Introduction

Genomic deep learning models that learn complex motifs and regulatory grammars from DNA sequence can predict molecular phenotypes such as transcription factor (TF) binding, histone modifications, chromatin accessibility, and gene expression directly from sequence [1, 2, 3, 4, 5, 6, 7, 8, 9]. These models have the potential to be powerful tools for analyzing personal genomes by predicting the molecular effects of individual genetic variants and haplotypes. In addition, interpreting the predictive sequence features learned by such models can provide biological insights into genome function and regulation. Approaches based on forward or backward propagation of influence can be used to identify DNA sequence motifs that drive model predictions [10]. However, because it can be difficult to fully interpret the learned sequence features and uncover mechanistic insights from “black box” neural networks, recent work has also aimed to develop explainable models that are more biologically interpretable [10, 11, 12]. For example, Puffin uses convolutional kernels of specific biologically-interpretable lengths to separate the contributions of simple sequence features from more complex binding motifs [11]. Here we develop a biologically-motivated approach to explicitly model the contributions of both DNA sequence and DNA biophysical or structural properties (DNA shape) to TF binding and other molecular phenotypes.

Shape readout, or the binding of a TF to a region of DNA based on its physical structure, is now considered a major mode of TF binding. Recent work increasingly examines local DNA shape features to explain TF binding outside of canonical sequence motifs [13, 14, 15]. An algorithm for finding shape motifs, or shape feature patterns that are enriched at TF binding sites, has also been proposed [16]. However, interpreting genomic deep learning models to extract their knowledge of how DNA shape affects TF binding remains underexplored. Deep neural networks capture extensive information about the mapping from genomic input to genomic activity, beyond what is captured by enrichment tests and other existing computational methods. Thus, understanding what is learned by a shape-based network can potentially shed insight on shape readout and reveal examples of shape-driven TF binding.

In this work, we present DeepShape, a deep convolutional neural network that incorporates DNA shape features alongside one-hot encoded DNA sequence as input, and apply this shape-aware modeling strategy to ChIP-seq peak prediction across 148 cell types and 170 TFs. We adopt two approaches to interpreting the trained DeepShape model, showing that feature attribution analyses based on the shape inputs enhance the interpretability of the model and provide additional insights into regulatory patterns not captured by sequence alone. Our results demonstrate that while sequence motifs are the primary drivers of TF binding, incorporating DNA shape features is useful for exploring non-canonical and co-bound sites.

## Materials and methods

### Model design

The deep convolutional neural network, DeepShape, was modeled after DeeperDeepSEA as implemented in the PyTorch based deep learning library Selene, which takes one-hot encoded DNA sequence as input [5]. Modifications were made to accommodate the addition of shape features. DeepShape processes 1000-bp sequence and shape inputs through two convolutional layers each, subsequently combining them to enable the model to capture unique information by learning separate representations for each input type. The combined representation passes through four additional convolutional layers, along with two max-pooling layers, three batch normalization layers, two dropout layers, and ReLU activations, before the output is flattened and passed through two fully connected layers, with the final output layer using a sigmoid activation function to predict the binary binding/not of each genomic target to the input sequence while considering the sequence’s DNA shape features (Fig. 1a). DeepShape uses a fixed 450 kernels per layer and filter length of 4, determined through a comprehensive hyperparameter search designed to adapt the model to the combined analysis of sequence and structural features (Supp. fig. 1).

**Figure 1:**
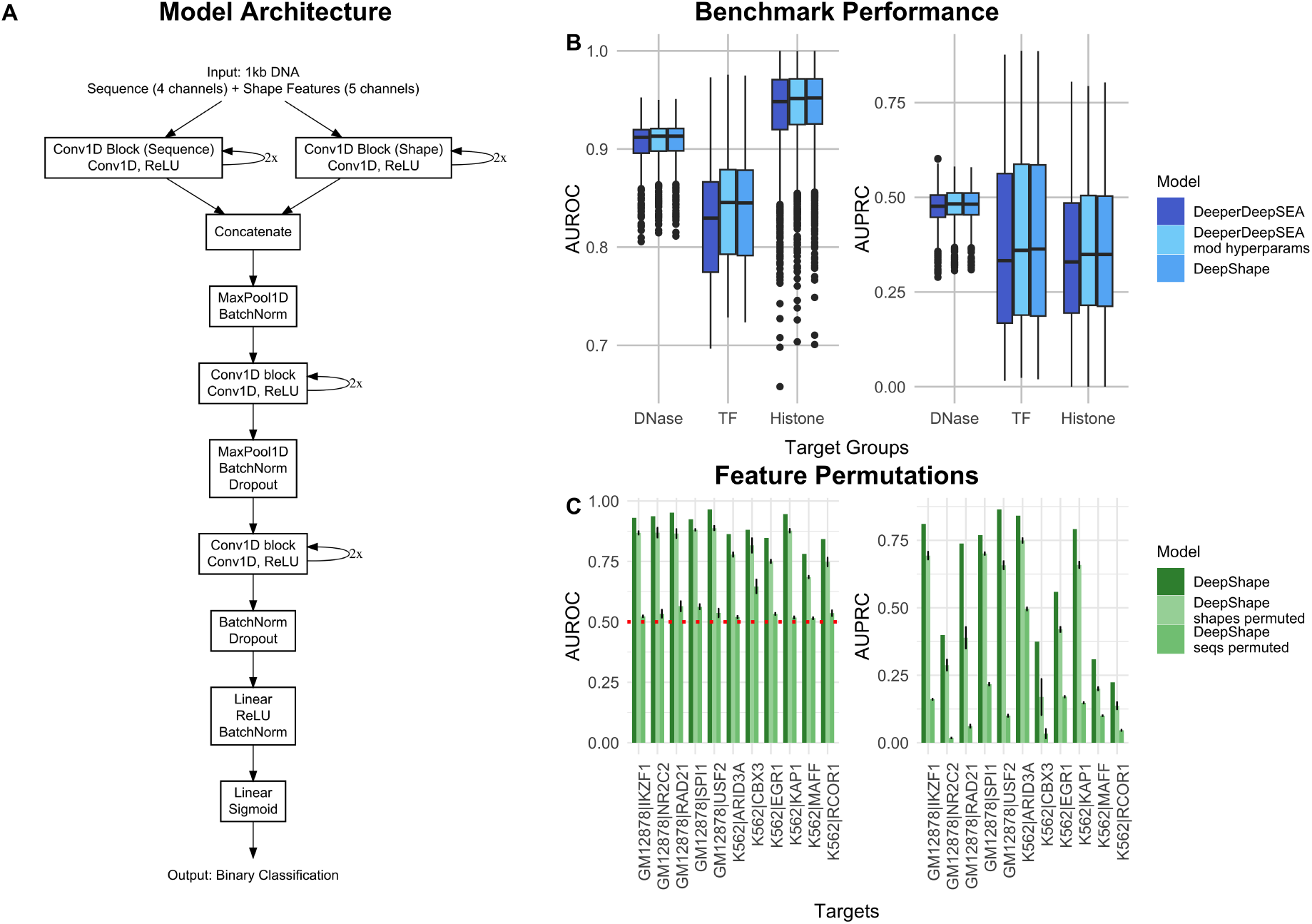
(a) Overview of the DeepShape model architecture. (b) Performance evaluation of DeepShape across held-out DNase, TF, and histone target groups. Modified hyperparameters improve sequence + shape and sequence only model. (c) Comparison of AUROC (left) and AUPRC (right) for 11 representative TFs (Data and training) based on trained DeepShape models from (b) versus models based on permuted shape or sequence features (bar colors). Permutations were performed for models with optimized hyperparameters.

To demonstrate this general approach, we implemented DeepShape to simultaneously predict the same 919 genomic tracks used to develop DeeperDeepSEA, each as a different output (“targets”). These include 690 TF targets (ChIP-seq), 125 open chromatin targets (DNase), and 104 histone mark targets (ChIP-seq) across 148 cell types and 34 treatment conditions. To create binary labels for these tracks, the genome was split into 200-bp bins and assigned a positive label if more than half the bin overlapped a peak region and was otherwise labeled negative, following the labeling strategy used in DeeperDeepSEA.

### Data and training

Training labels were computed from uniformly processed ENCODE and Roadmap Epigenomics data releases. The 11 TF targets used for hyperparameter tuning were selected based on their strong shape or sequence preferences, respectively, as described by Samee et al. (2019) [16]. The 919 chromatin features used to benchmark performance against DeeperDeepSEA were those used by the original DeepSEA and DeeperDeepSEA model evaluations. We set the training parameters (number of steps, batch size, and related configurations) to match how both DeepSEA and DeeperDeepSEA were originally trained [1, 5]. The benchmark was performed using the following reciprocal validation/test splits (six total): chromosomes 6 and 7, 8 and 9; 9, and 10, 11 and 12; 14 and 17, 15 and 16. See Code and data availability for configuration files detailing the training set up.

### Shape inputs

Whole genome shape features were generated for five structural attributes of DNA—minor groove width, helical twist, propeller twist, and electrostatic potential—that have been found to be among the most important for evaluating protein-DNA readout modes [17]. The set of five shape features was generated by passing the hg19 reference FASTA to DNAShapeR, a software package that uses a sliding pentamer window to generate shape features from all-atom Monte Carlo simulations [18]. The output of this provides a single-nucleotide resolution representation of DNA conformation within local contexts, with the predicted structural outputs representing preferred conformations that are intrinsic to a given DNA sequence [19]. Gaps in the resulting shape assemblies were filled with the mean shape feature value for each of the five shape features.

### DeepLIFT

To quantify the contribution of each input feature to the prediction of TF binding, we employed DeepLIFT (Deep Learning Important FeaTures) [20]. For a given TF track, DeepLIFT assigns an attribution score to each feature, indicating its influence on the model’s prediction. These attributions are computed such that they satisfy the summation-to-delta property:

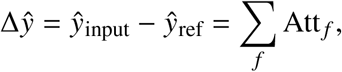

where *ŷ*_input_ represents the model’s prediction for the given input sequence, *ŷ*_ref_ is the prediction for a reference sequence, and Att _*f*_ denotes the attribution for feature *f*. The sum of all feature attributions equals the difference between the prediction for the input sequence and that for the reference.

Since the model predicts a binding probability in the range [0, 1] for each TF track, and the binding probability for a random sequence typically approaches zero, Δ*ŷ* approximates the predicted probability *ŷ*_input_.

## Results

### DeepShape model overview

DeepShape extends the DeeperDeepSEA architecture by incorporating DNA shape features alongside sequence inputs (Methods). After training on the full set of targets across six distinct chromosome splits (Methods), we benchmarked the performance of DeepShape (sequence + shape) against a DeeperDeepSEA model that we trained (sequence-only).

We find that DeepShape has similar or slightly improved performance relative to DeeperDeepSEA on the three types of prediction tasks (Fig. 1b), with an average AUROC of 0.929 and average AUPRC of 0.379 compared to DeeperDeepSEA’s average AUROC of 0.925 and average AUPRC of 0.367. The performance of the DeeperDeepSEA model we trained was similar to that of the published model. Our focus was on TF binding, but we included open chromatin and histone targets because these genomic elements can be bound by TFs and hence predicting them alongside the TF targets may improve performance. Using all 919 tracks also enabled us to directly compare results with the published DeeperDeepSEA model. We observed that modifying DeeperDeepSEA’s hyperparameters to match those of DeepShape also improved its performance relative to the base-line DeeperDeepSEA model (Fig. 1b), suggesting that the hyperparameter tuning improved model representations of sequence inputs effectively as well.

To evaluate the degree to which DeepShape leverages shape information for predicting TF binding, we permuted shape features across each batch by setting the shape features of the *i*th example to the shape features of the *σ*(*i*) example, where *σ* is a random permutation (Fig. 1c). We separately permuted sequence features for comparison. This permutation analysis resulted in an average AUROC contribution ratio of 0.228 and average AUPRC contribution ratio of 0.348 for shape versus sequence contributions in DeepShape, indicating that sequence information is most important for predicting TF binding, as expected, but that shape also contributes notably to model predictions. A similar analysis of a sequence-and-shape model with the same original hyperparameters (filter size, number of kernels per layer) as DeeperDeepSEA shows lower shape contributions (average AUROC contribution ratio = 0.101, average AUPRC contribution ratio = 0.269; Supp. fig. 2). This suggests that the optimized hyperparameters enable better use of shape inputs.

**Figure 2:**
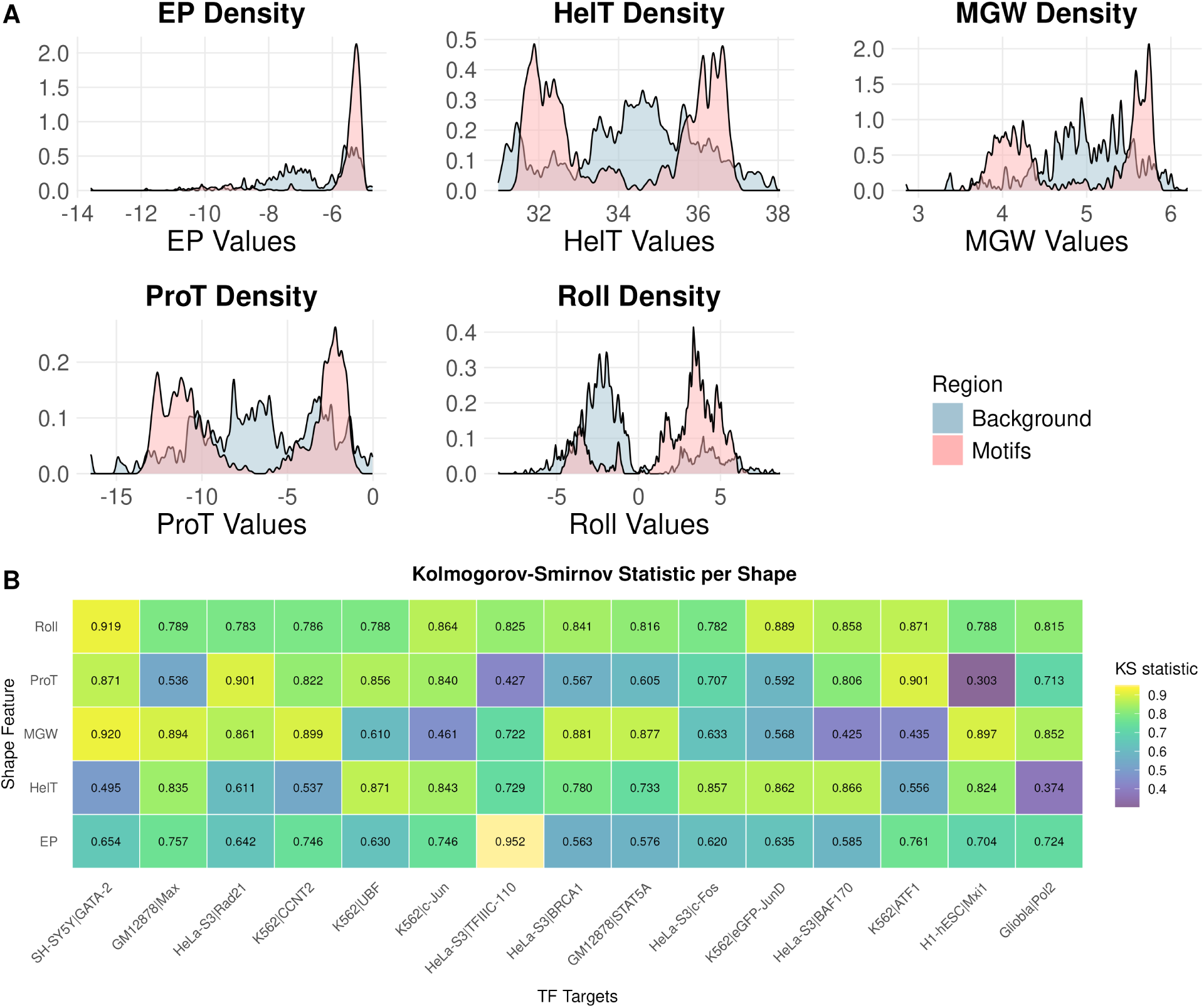
Distribution of DNA shape feature values at positions with high attribution scores. (a) Density plots compare background distributions to positions with high DeepLIFT attribution score. High-attribution positions refer to nucleotide positions within motifs that exhibit high average DeepLIFT attribution across aligned seqlets. Positions with high attribution scores exhibit a shift in value distributions relative to the background, suggesting structural patterns associated with regulatory activity. The x-axis represents the specific range of values for each shape feature. (b) Kolmogorov-Smirnov (KS) statistics quantifying the degree of separation between motif and background distributions for each shape feature across 15 TF targets. Higher KS values indicate stronger motif-background divergence. Variation across both features and targets indicates that shape contributions are target-specific.

### Model interpretability

Although the joint sequence-and-shape model performs similarly to the sequence-only model with the modified hyperparameters, we hypothesized that DeepShape’s explicit modeling of shape inputs would provide greater interpretability. To better understand the contributions of sequence and DNA shape features to TF binding predictions, we performed a series of interpretability analyses. These included activation-based analyses to examine how specific features influence model predictions and attribution-based approaches (DeepLIFT) to identify high-contribution positions across input features (Methods) [20]. Using these attributions, we investigated patterns in sequence and shape features that are enriched at binding sites.

Compared to background sequence distributions, shape values in bound regions revealed distinct shifts, suggesting that structural features play a key role at regulatory sites (Fig. 2a). To examine variation across individual TF targets, we computed Kolmogorov-Smirnov (KS) statistics for each shape feature and find variation in the degree of motif-background separation across both features and targets (Fig. 2b), suggesting that individual targets may exhibit preferences for specific shape features, consistent with observations from previous studies [16]. These patterns are reflected in individual TF targets. For example, the distribution of MGW values in GM12878|MEF2A shows a strong shift toward narrower minor groove widths in motif regions (Supp. fig. 3a), consistent with prior structural studies showing that MEF2A recognizes its binding site by selecting a narrow minor groove width and engaging two base pairs per half-site [21]. In contrast, K562|CEBPB displays a shift in Roll values, with motif regions enriched for higher Roll compared to background (Supp. Fig. 3b). This differs from previous shape motif analyses, which identified HelT as the preferred feature for CEBPB [16], suggesting that Roll may also play a role in CEBPB binding, depending on cellular context or motif instance.

**Figure 3:**
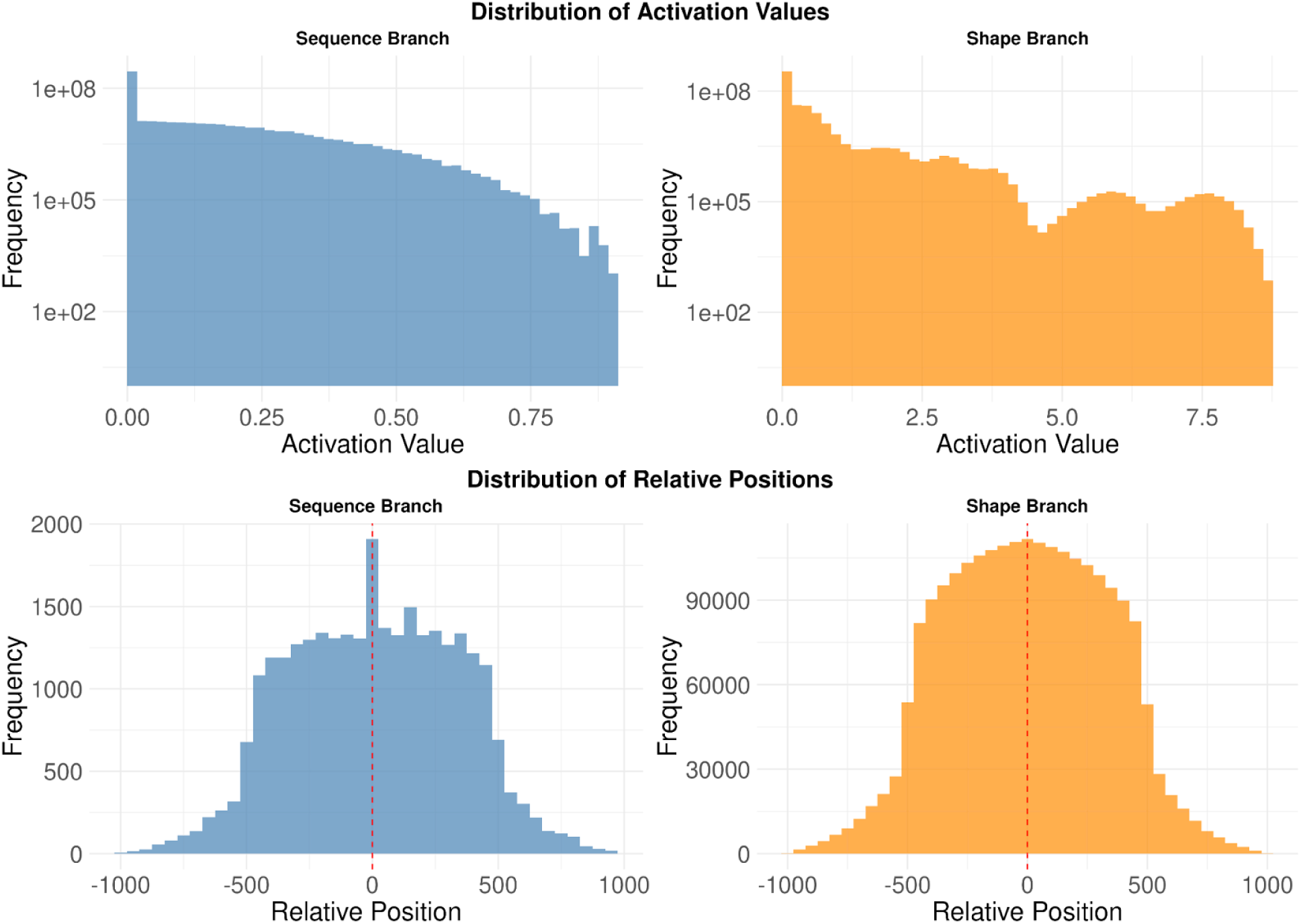
Distribution of activation values across filters in the sequence (top left) and shape (top right) branch. Distribution of relative positions of activations above a given threshold in the sequence (bottom left) and shape (bottom right) branch. Sequences containing a single CACGTG motif are aligned such that the motif is at position 0.

To gain additional insight into feature importance, we compared the average number of high-attribution positions per motif type (Supp. table 1). Sequence motifs were associated with a higher number of high-attribution positions compared to shape motifs, reflecting their role in capturing canonical motifs directly involved in TF binding, while shape motifs highlighted shorter segments, providing complementary structural information. These analyses demonstrate that sequence and shape features provide distinct yet interrelated insights into regulatory predictions.

These findings raised the question of how sequence and shape features contribute in specific binding contexts. To explore this, we applied DeepShape to the Myc-Max TF heterodimer, examining how sequence and DNA shape features contribute to binding specificity. *In vivo* studies show that Max binds E-box motifs in approximately five times as many locations as Myc. Prior work identified distinct shape motifs in Myc-Max ChIP-seq peaks compared to Myc-unbound-Max regions, suggesting that DNA shape may play a role in their differential binding patterns [16]. We trained DeepShape on a single Myc-Max track based on the intersection of Myc and Max ChIP-seq peaks, 3,156 peaks in total, in GM12878 cells. Among these, 902 peaks contained exactly one CACGTG motif (E-box), while 1,071 peaks contained one or more CACGTG motifs. The sequence and shape branches were evaluated separately by applying their filters and obtaining activations for Myc-Max peaks containing a single CACGTG motif. The model’s sequence layer exhibits a sharp peak in the distribution of relative positions centered around the E-box motif. In contrast, the shape branch displays a more diffuse distribution of activations without sharp peaks and with elevated values flanking rather than overlapping the E-box (Fig. 3).

This activation pattern is consistent with prior studies showing that while the E-box motif plays a central role in Myc-Max binding, it is not sufficient to fully explain specificity. Guo et al. (2014) demonstrated that Myc binding frequently depends on chromatin context and transcriptional machinery rather than sequence motifs alone [22]. Allevato et al. (2017) further showed that Myc-Max binds both canonical and non-canonical motifs, with flanking sequences and lower-affinity binding sites modulating interactions [23]. Additionally, Yang et al. (2014) highlighted the importance of DNA shape features in refining binding predictions, particularly in regions adjacent to sequence motifs [24]. These findings support the idea that DNA shape features captured by the model’s shape branch may provide supplementary information for fine-tuning of binding specificity.

To assess the relative contributions of sequence and shape features to binding predictions across different TF targets, we compared their attribution values (Fig. 4a). Across most TFs, we observed a consistent relationship between sequence and shape attributions: shape attribution is generally 0.269 times the sequence attribution, as indicated by the line of best fit, though TFs show variability around this trend. This suggests that while sequence features contribute more substantially to binding predictions overall, shape features provide a supplementary contribution across diverse TFs with some TFs relying on shape features more than others.

**Figure 4:**
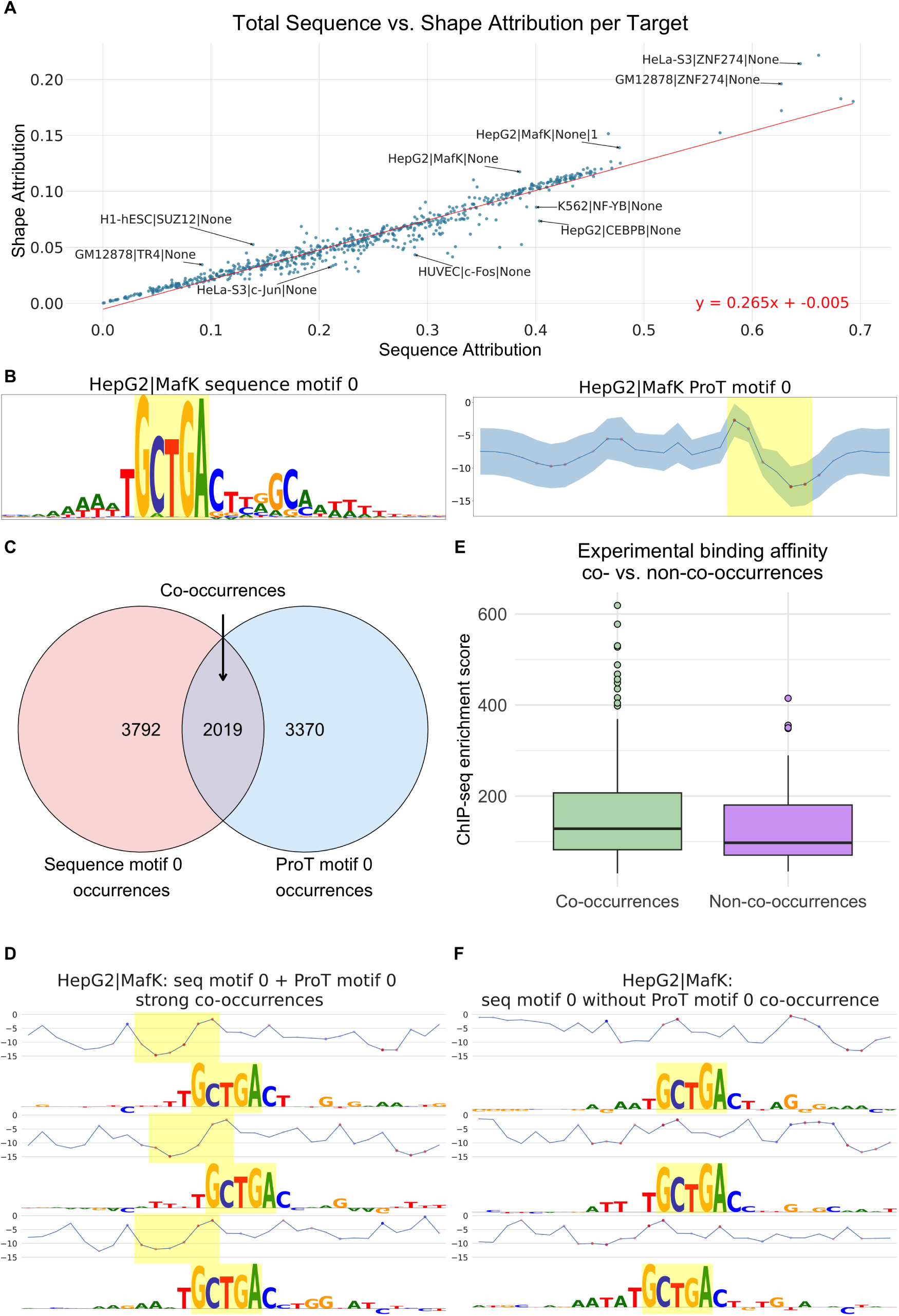
Sequence vs. shape attributions and co-occurrence analysis of HepG2|MafK motifs. (a) Total sequence versus shape attribution for each TF target. Each point represents a TF target, with attributions averaged across bound sequences. The best fit line in red indicates the general trend, with notable deviations labeled. (b) Sequence motif logo (left) and average profile with spread for ProT motif (right) for HepG2|MafK. High-attribution regions of each motif are highlighted. (c) Venn diagram showing the number of co-occurrences of sequence motif 0, ProT motif 0, and their co-occurrences. (d) Examples of strong co-occurrences between sequence motif 0 and ProT motif 0 in HepG2|MafK. The ProT profile is shown above the sequence logo for each example, with high-attribution regions highlighted. (e) ChIP-seq enrichment scores for strong sequence motif 0 occurrences that co-occur with ProT motif 0 versus those that do not. Co-occurrences exhibit significantly higher enrichment (*p* = 8.7 × 10^−4^). (f) Examples of sequence motif 0 occurrences without co-occurrences of ProT motif 0.

Among the TFs analyzed, MafK stood out as having relatively higher shape attributions compared to sequence attributions (Fig. 4a), indicating that DNA shape features might play a unique role in modulating MafK binding specificity. To further investigate this, we used DeepLIFT to score each input feature based on its contribution to the prediction of the network, and then ran TF-MoDISco to uncover sequence and shape motifs learned by the model (Methods). TF-MoDISco takes a set of genomic sequences and their attributions as input, and it clusters high-attributions subsequences called “seqlets” to generate motifs [25]. We adapted TF-MoDISco to run on shape feature attributions by transforming each quantitative shape feature into its quartile, producing a 4-dimensional one-hot encoding. As a result, for any type of shape feature, we can obtain shape motifs that explain the binding of a particular TF target. For the HepG2|MafK target, TF-MoDISco identified a sequence motif matching the canonical TGCTGA(C/G)TCAGCA MafK motif, along with a propeller twist (ProT) shape motif (Fig. 4b). The ProT motif is characterized by a peak followed by a pronounced dip in the high-attribution region, which is not in the canonical motif’s profile.

We examined the co-occurrence of these motifs, defining an occurrence as a seqlet belonging to the corresponding TF-MoDISco motif cluster. The ProT motif frequently co-occurs with the sequence motif, typically at the end of the sequence motif and in the nearby flanking region (Fig. 4c, d). To assess the potential modulatory effect of the ProT motif on binding affinity, we compared ChIP-seq signals for strong sequence motif occurrences (> 90th percentile Pearson similarity of attributions) that co-occur with the ProT motif versus those that do not. Strong sequence motif occurrences co-occurring with the ProT motif showed significantly higher binding (Fig. 4e, Supp. table 2).

Examination of individual occurrences revealed that the downward ProT dip often corresponds to A-tracts or T-tracts flanking the sequence motif (Fig. 4d, f). This finding aligns with previous observations that A-tracts and T-tracts significantly affect DNA shape [26]. These results suggest that DNA shape features in flanking regions can modulate binding to canonical sequence motifs, potentially explaining some of the binding specificity not captured by sequence alone.

After identifying sequence and shape motifs using TF-MoDISco, we evaluated their relative contributions to TF binding predictions. To do this, we used the motifs to featurize our dataset and trained logistic regression and gradient boosting models to predict TF binding (Supp. fig. 3). Each sequence was featurized by computing the maximum motif match score across the sequence for each motif, resulting in an n-dimensional representation where n is the number of motifs, performed separately for each target. For model training, five sequence motifs were selected using forward step-wise selection with logistic regression and cross-validation, and five shape motifs were selected in the same way. These 10 motifs were then used to train both logistic regression and gradient boosting classifiers. Across targets, the mean ± standard deviation number of motifs from which these 10 were selected was 21 ± 9 for sequence motifs and 114 ± 26 for shape motifs.

Our first conclusion based on these results is that sequence motifs are the primary drivers of TF binding predictions. When examining the most important motif for each assay, as determined by logistic regression coefficient significance and gradient boosting feature importance, we found that sequence motifs were deemed most important in 95.3% and 82.9% of cases, respectively (Fig. 5a). This suggests that while shape features contribute to the models’ performance, sequence information remains the dominant factor in predicting TF binding.

**Figure 5:**
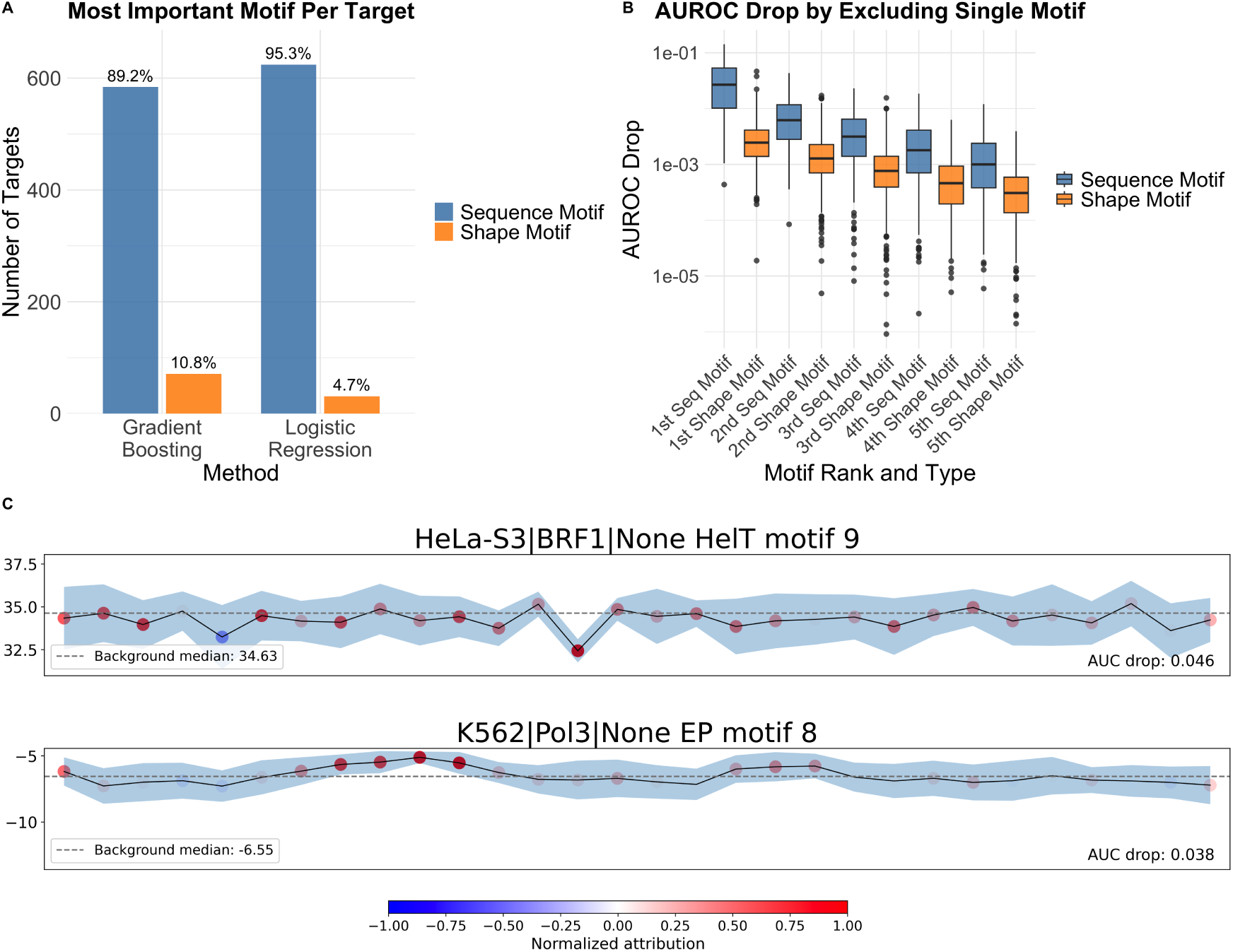
Motif importance and activation patterns across sequence and shape features in TF binding. (a) Percentage of TF targets where, out of the 10 selected motifs, a sequence / shape motif was deemed most important, according to logistic regression coefficient significance and gradient boosting feature importance. (b) Performance drops resulting from ablating each individual motif and retraining the gradient boosting model. “*i*th motif” refers to the motif for that TF target which resulted in the *i*th highest performance drop measured via AUROC. (c) Shape motifs from two representative TF targets—HeLa-S3|BRF1 and K562|Pol3—that exhibited the largest AUROC drops upon ablation of a shape motif, as shown in (a).

To further quantify the relative contributions of sequence and shape motifs, we performed ablations by removing individual motifs and retraining our gradient boosting model (Fig. 5b). The performance drops resulting from feature removal confirm that for most TFs there is a single sequence motif that has a much larger impact on the model than any other sequence or shape motif. Nonetheless, the model remained fairly performant after dropping the top motif, indicating that DeepShape has learned more than simply canonical sequence motifs. These findings confirm the primacy of sequence information in determining TF binding specificity, while underscoring that shape motifs and non-canonical sequence motifs also contribute to predictions across TFs. For certain targets, shape motifs were among the top contributors, with their removal leading to measurable performance drops (Fig. 5c), further supporting their complementary role in binding prediction.

## Discussion

DeepShape’s ability to integrate both sequence and shape information provides a more comprehensive understanding of DNA-protein interactions. Our findings demonstrate that while sequence motifs are the primary drivers of TF binding, DNA shape features play a role in modulating this binding.

The uniformity of the shape to sequence attribution ratio across TF targets is noteworthy, and may reflect the model’s consistent utilization of shape information rather than variations in biophysical shape readout mechanisms among TFs. Targets deviating from this trend, such as HepG2|MafK, could potentially indicate TFs with more pronounced shape-dependent binding mechanisms.

The example of HepG2|MafK binding illustrates how shape motifs, particularly in combination with sequence motifs, can fine-tune binding affinity. The propeller twist shape motif identified in this case suggests that local DNA structural properties in flanking regions can significantly influence binding to canonical sequence motifs. This observation aligns with the emerging view of shape readouts as an important mechanism in TF-DNA recognition.

Our approach has notable limitations. The DNA shape features used represent a static, localized view of DNA structure, neglecting its dynamic nature and larger-scale features like topologically associated domains, or deviations in shape due to modifications such as methylation. Additionally the use of continuous values for shape inputs may contribute to the observed diffuse attribution patterns. This could be tested by converting input features to a binary form, such as categorizing them into extreme versus non-extreme quantiles. Further, future research could explore incorporating shape features as an output track, aiding in interpreting what the model has learned about structure directly from DNA sequence.

Despite these limitations, our study demonstrates the value of integrating DNA shape information into genomic deep learning models. DeepShape provides an additional axis for model interpretability, revealing regulatory patterns not evident from sequence alone. This approach offers insights into cases where sequence motifs incompletely explain binding patterns.

## Code and data availability

Code to train and evaluate DeepShape models and run TF-MoDISco interpretability pipelines are available at https://github.com/ni-lab/DeepShape. DNA shape inputs are available on Zenodo (DOI 10.5281/zenodo.13334119). Model weights are available on Zenodo and https://huggingface.co/ryankeivanfar/DeepShape.

## Acknowledgements

We thank Shiron Drusinsky, Sean Whalen, Jodi Lee, and other members of the Pollard and Ioannidis labs for helpful discussions. This work was supported by the UC Noyce Initiative for Computational Transformation, Additional Ventures, Gladstone Institutes, the W. M. Keck Foundation and the Biswas Family Foundation. We used the Savio computational cluster resource provided by the Berkeley Research computing program at the University of California, Berkeley (supported by the UC Berkeley Chancellor, Vice Chancellor for Research, and Chief Information Officer). N. M. I. and K. S. P. are Chan Zuckerburg Biohub San Francisco Investigators.

## Supplementary material

**Supplementary figure 1:**
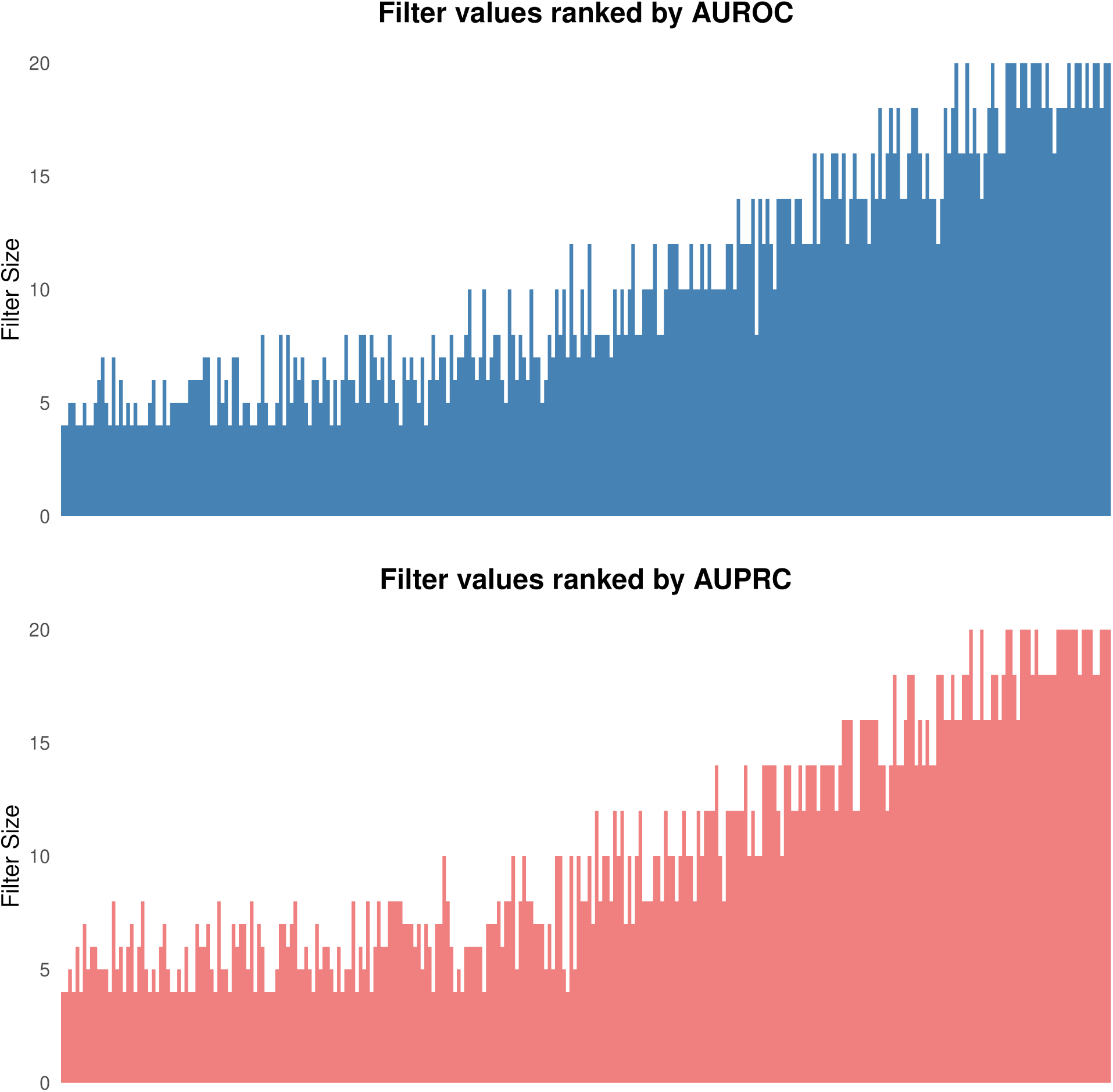
Performance trends from a hyperparameter search on a version of DeepShape that used only DNA shape features as input. Filter sizes and number of filters per layer were tested to optimize shape feature capture for predicting TF binding. Smaller filter sizes demonstrated a trend toward improved performance. Filter sizes ranked by AUROC (top) and AUPRC (bottom), sorted from highest to lowest.

**Supplementary figure 2:**
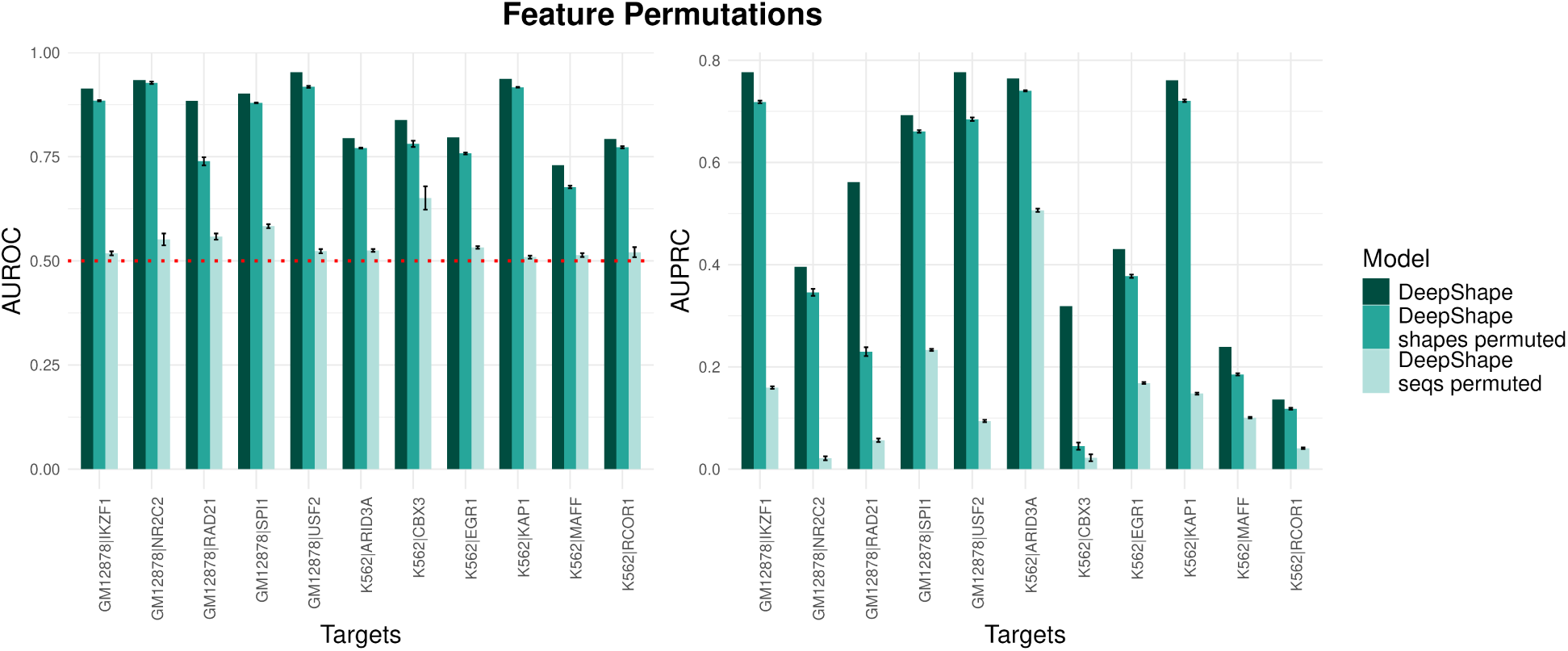
Comparison of AUROC (left) and AUPRC (right) for 11 representative TFs (Data and training) based on a trained DeepShape model versus models based on permuted shape or sequence features (bar colors). Permutations were performed with the same hyperparameters as the published DeeperDeepSEA model.

**Supplementary figure 3:**
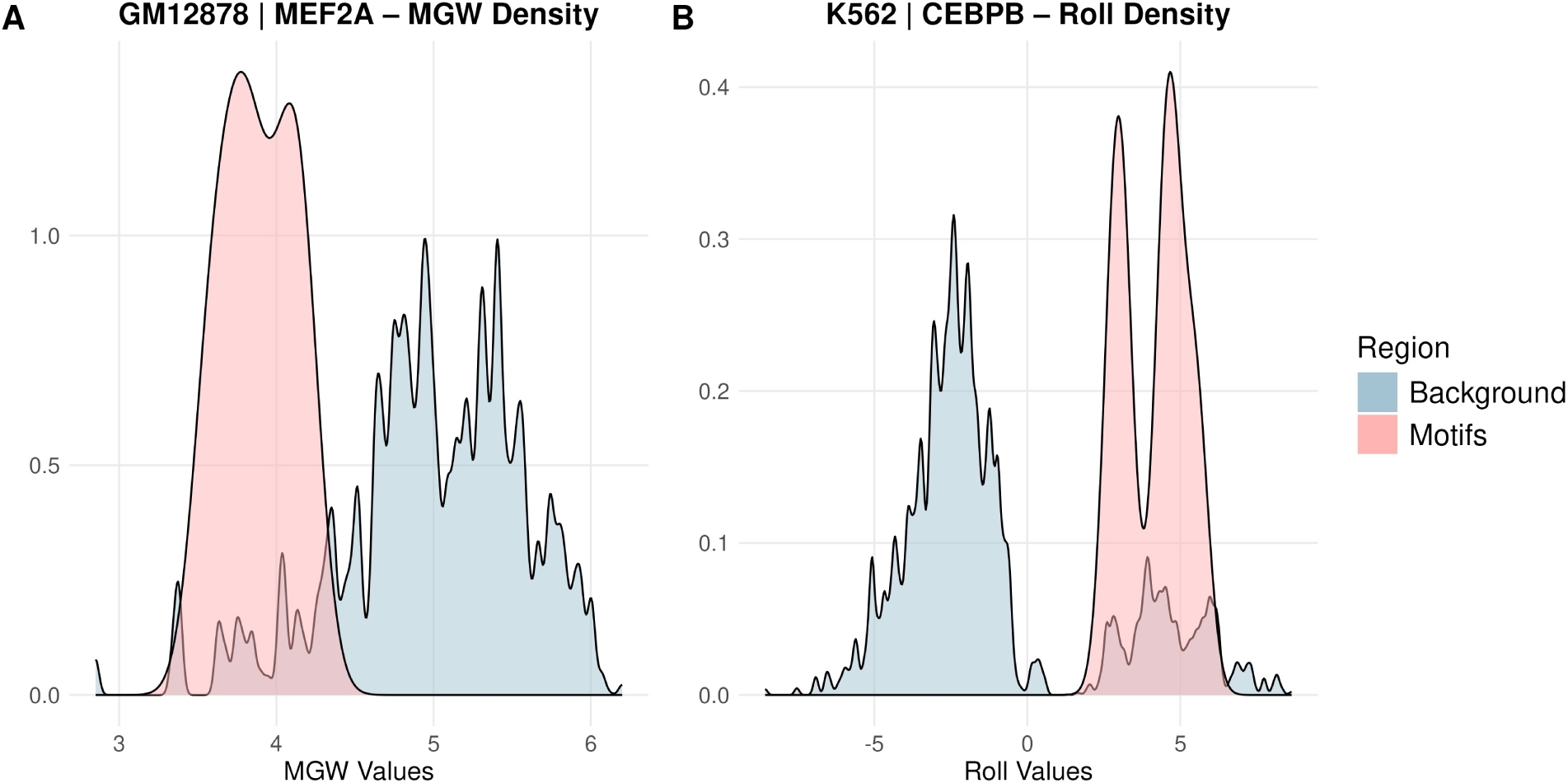
(a) Minor groove width (MGW) distributions for GM12878|MEF2A, showing a preference for narrower minor groove widths relative to background. (b) Roll distributions for K562|CEBPB, showing a shift toward higher roll values in motif regions compared to background.

**Supplementary figure 4:**
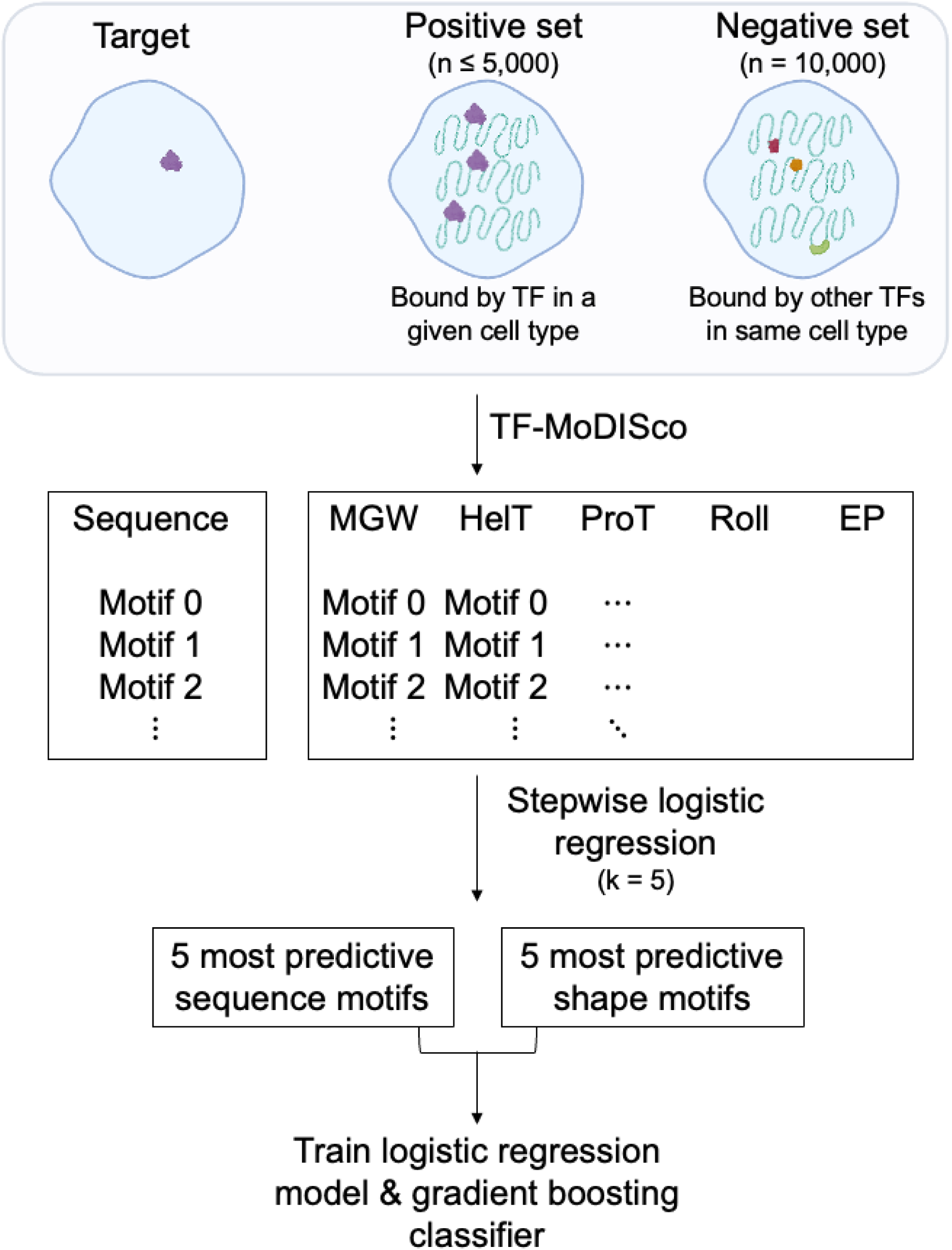
Pipeline for identifying predictive sequence and shape motifs. Positive sets included sequences bound by the TF of interest, while negative sets included sequences bound by other TFs in the same cell type. TF-MoDISco extracted sequence and shape (MGW, HelT, ProT, Roll, EP) motifs from DeepLIFT attribution scores. The most predictive motifs were ranked using stepwise logistic regression, with the top five sequence and shape motifs used to train logistic regression and gradient boosting models, highlighting the contributions of sequence and shape features to TF binding predictions.

**Supplementary table 1:** Average number of high-attribution positions per motif for sequence features and each DNA shape feature. Sequence motifs consistently exhibit a greater number of high-attribution positions.

| **Motif type** | **Avg. no. high-attribution positions per motif** |
| --- | --- |
| Sequence | 5.46 |
| MGW | 3.02 |
| HelT | 3.35 |
| ProT | 3.19 |
| Roll | 3.37 |
| EP | 3.72 |

**Supplementary table 2:** Summary statistics of co-occurring and non-co-occurring strong sequence motif occurrences in HepG2|MafK. Co-occurring sequence motifs exhibit significantly higher ChIP-seq enrichment scores compared to non-co-occurring motifs.

| **Metric** | **Value** |
| --- | --- |
| No. co-occurring strong sequence occurrences | 353 |
| No. non-co-occurring strong sequence occurrences | 170 |
| Mean ChIP-seq score of co-occurrences | 157.5 |
| Mean ChIP-seq score of non co-occurrences | 129.8 |
| One-sided p-value | 8.7e-04 |

